# Intranasal HSV-1 Infection Drives Region-Specific Interferon-Dominant Microglial Remodeling

**DOI:** 10.64898/2026.03.13.711627

**Authors:** Seth Frietze, Cameron Lunn, Dean Oldham, Joseph R Boyd, Andrew N. Bubak, Amalia Bustillos Saucedo, Maria A Nagel, Diego Restrepo, Kimberley D Bruce, Christy S Niemeyer

**Author notes:** Corresponding authors: Kimberley D Bruce, Christy S. Niemeyer.

## Abstract

**Background and Objectives:** Herpes simplex virus type 1 (HSV-1) is a neurotropic pathogen capable of invading the central nervous system (CNS) and increasingly associated with chronic neuroinflammation, cognitive impairment, and neurodegenerative disease. While microglia orchestrate the initial immune response to HSV-1, the molecular mechanisms that regulate their sustained neuroinflammatory activity *in vivo* remain poorly understood.

**Methods:** To define the transcriptional and epigenetic mechanisms that shape microglial responses during acute HSV-1 infection *in vivo,* we have, for the first time, integrated single-nucleus RNA sequencing, chromatin accessibility profiling, and spatial transcriptomics in a physiologically relevant intranasal HSV-1 infection model.

**Results:** Single-cell multiome analysis of CD11b⁺ nuclei identified transcriptionally and epigenetically distinct microglial and macrophage populations. HSV-1 infection redistributed monocyte-lineage states, with a marked overrepresentation of interferon (IFN)-responsive microglia and macrophage-associated populations. These states exhibited amplification of STAT1/2-, IRF1-, and CEBPB-centered regulons, distinguishing IFN-responsive microglia from macrophage-enriched populations rather than reflecting uniform activation. Homeostatic microglial gene signatures (*e.g., ApoE, Cst3*) were reduced in response to HSV-1 infection. Spatial transcriptomics localized HSV-1 antigen to discrete brainstem regions, which were enriched for predicted STAT-, IRF-, and CEBPB-regulated targets identified through single-nuclei analysis.

**Discussion:** Using a multiomic framework, we demonstrate that HSV-1 infection drives transcriptional and epigenetic remodeling of microglial populations, characterized by a dominance of IFN-responsive states and a loss of homeostatic signatures. These findings provide mechanistic insight into how localized viral infection can reprogram microglial regulatory landscapes to maintain persistent HSV-1–associated neuroinflammation, contributing to long-term neurological vulnerability and neurodegenerative disease risk.

## Introduction

Herpes simplex virus type 1 (HSV-1) is a neurotropic alphaherpesvirus that establishes lifelong latency in peripheral sensory ganglia and can access the central nervous system (CNS) via retrograde transport along trigeminal or olfactory pathways^1^. While HSV-1 is best known for causing recurrent mucosal lesions (‘cold sores’), it can also cause rare viral encephalitis and has increasingly been implicated in chronic neuroinflammation and long-term cognitive impairment, including Alzheimer’s disease and related dementias ^2–8^. Since HSV-1 can access the CNS through anatomically defined neural routes, this raises the possibility that localized innate immune responses may shape long-term vulnerability in specific brain regions. Understanding how antiviral immune responses reshape microglial function may provide insights into mechanisms linking viral infection to long-term neurological vulnerability.

Innate antiviral responses within the CNS are coordinated by resident microglia and recruited macrophages^9–12^. These cells function as immune sentinels and maintain tissue homeostasis through surveillance, synaptic remodeling, lipid metabolism, and phagocytosis^13–18^. Microglia exhibit substantial transcriptional and functional plasticity, rapidly responding to internal and external triggers. In response to viral infection, microglial populations adopt diverse transcriptional states associated with inflammatory signaling, antigen presentation, and antiviral defense^11, 19–23^. While microglial plasticity is an essential adaptive response^24^, maladaptive remodeling of persistent microglial states can predispose neuroinflammatory and neurodegenerative disease^25^. Yet whether similar remodeling occurs following HSV-1 infection remains largely unknown.

In murine models, intranasal infection results in focal viral antigen within discrete brainstem regions, providing a tractable system to examine regionally confined antiviral responses^1^. During HSV-1 infection, microglia rapidly respond, surrounding infected neurons and astrocytes and exhibiting a swollen yet ramified morphology^10, 11^. Both *in vitro* and *in vivo* studies have also shown that in response to HSV-1 infection, microglia increase the production of pro-inflammatory cytokines and overexpress markers of activation^9–13^. In addition, *in vitro* studies have shown that HSV-1 infection reprograms microglia away from a phagocytic phenotype^36^ and downregulates the expression of phagocytic markers^37^. These functional shifts are notable because microglial programs are critical for maintaining neuronal health and clearing pathological protein aggregates associated with neurodegenerative disease^26^. While previous studies highlight functional remodeling of microglia during HSV-1 infection, the regulatory mechanisms that govern the emergence and persistence of these states in chronic or neurodegenerative diseases remain unknown.

The engagement of microglia states is coordinated by transcription factor–centered gene regulatory networks that link inflammatory cues to chromatin accessibility and downstream gene expression programs^27–29^. Growing evidence suggests that inflammatory stimuli can induce epigenetic remodeling in microglia, establishing transcriptional programs that persist beyond the acute immune response and contribute to chronic neuroinflammation and neurodegeneration^30, 31^. Defining how epigenetic remodeling is engaged at peak activation is essential for understanding how disease-associated microglia states are established during HSV-1 infection. However, how distinct microglial populations deploy these regulatory programs during acute HSV-1 infection *in vivo* remains incompletely defined.

Recent single-cell and spatial transcriptomic studies have revealed previously unrecognized heterogeneity in antiviral responses within the infected brain^29^. These analyses have identified a virus-associated microglial population enriched for type I interferon (IFN-I) signaling, which emerges alongside infiltrating monocytes and T cells and contributes to broader transcriptional remodeling across multiple cell types within the CNS. However, these studies primarily characterize transcriptional states and do not resolve the underlying regulatory architecture, including chromatin accessibility and transcription factor networks, that drive persistent IFN-dominant microglial responses. Given persistent IFN signaling has been linked to pathological microglial activation^32–34^. How IFN-associated regulatory programs shape microglial populations during HSV-1 infection is key to understanding how antiviral responses reshape microglial identity and prime the CNS for long-term neuroinflammatory or neurodegenerative outcomes.

To address these questions, we profiled CD11b^+^ microglia isolated at 6 days post–intranasal HSV-1 infection, a stage marked by robust innate immune HSV-1 activation^1^. For the first time, we integrated single-nuclei transcriptomics, chromatin accessibility profiling, and spatial transcriptomics to identify the regulatory architecture of antiviral microglial states during acute HSV-1 infection. This multimodal approach revealed the emergence of IFN-dominant microglial subpopulations characterized by STAT1/2-, IRF1-, and CEBPB-centered transcription factor regulons following HSV-1 infection. Notably, these regulatory programs denote transcriptional changes within regions containing viral antigen, characterized by elevated IFN signaling and reduced homeostatic signatures. Together, these analyses define the transcriptional and epigenetic mechanisms regulating HSV-1–associated microglia states and resolve single-cell regulatory programs within spatially defined anatomical boundaries.

## Results

### Single-nuclei multiome and spatial transcriptomic profiling of acute HSV-1 infection

To define how microglial populations respond to acute HSV-1 infection, we combined single-nuclei multiome profiling with spatial transcriptomics (**Fig. 1A**). CD11b⁺ nuclei were isolated from whole brains for paired single-nuclei RNA sequencing (snRNA-seq) and single-nuclei assay for transposase-accessible chromatin sequencing (snATAC-seq). In a separate cohort subjected to the same infection protocol and collection time point, brain sections were analyzed using GeoMx Digital Spatial Profiling (DSP) with immunofluorescent labeling for HSV-1 antigen (gB) and Iba1 to guide region-of-interest selection.

**Figure 1.**
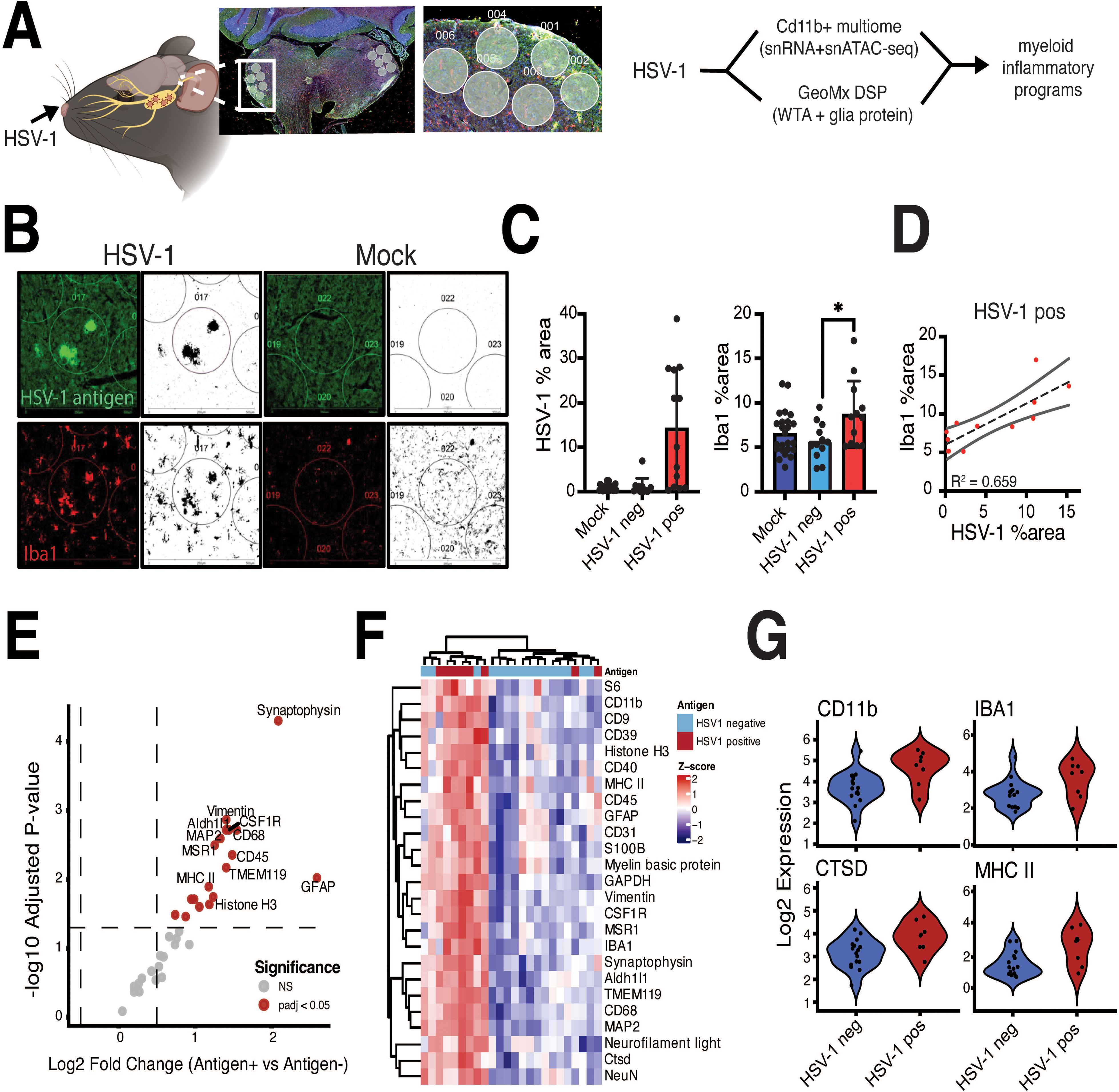
Spatially restricted HSV-1 infection and microglial response in the trigeminal nucleus of the brainstem. **A)** Adult C57BL/6J mice were intranasally infected with non-lethal HSV-1 (Makrae strain) or mock-treated and analyzed at 6 days post-infection. CD11b⁺ nuclei were isolated from whole brains for paired single-nucleus RNA-seq (snRNA-seq) and single-nucleus ATAC-seq (snATAC-seq). In a separate cohort subjected to the same infection protocol, brain sections were processed using the GeoMx Digital Spatial Profiler (DSP) with immunofluorescent labeling for HSV-1 antigen (gB) and Iba1 to guide region-of-interest (ROI) selection. Representative whole-section image from HSV-1 infection and ROI masks are shown. **B)** Immunofluorescence images from HSV-1-infected and mock-treated brains. HSV-1 antigen (green) localizes predominantly to the ipsilateral spinal trigeminal nucleus (SpVN) in infected animals and is absent in mock controls. Iba1 (red) signal is increased in antigen-positive regions. Grayscale images show segmentation masks used for ROI selection. **C)** Quantification of HSV-1 antigen-positive area (left) and Iba1-positive area (right) across mock, HSV-1 antigen-negative, and HSV-1 antigen-positive regions. Bars represent mean s.e.m.; points represent individual regions. **D)** Correlation between HSV-1 antigen area and Iba1-positive area in HSV-1-positive regions. Each point represents a single region; line indicates linear regression with 95% confidence interval. **E)** Differential protein enrichment analysis across the DSP glial protein panel comparing HSV-1 antigen-positive and antigen-negative ROIs. Protein abundance values were normalized and globally centered across ROIs prior to linear modeling. Proteins with adjusted P value < 0.05 (Benjamini-Hochberg FDR) are highlighted in red; non-significant proteins are shown in gray. **(F)** Per-protein Z-score–scaled normalized DSP protein abundance values across HSV-1 antigen–positive and antigen–negative ROIs. **(G)** Log₂-normalized DSP protein abundance values for selected markers (CD11b, IBA1, CTSD, MHC II) across HSV-1 antigen-negative and antigen-positive ROIs. CD11b *p =* 0.0159; IBA1 *p =* 0.0926; CTSD *p =* 0.023; MHC II *p =* 0.00875

Adult C57BL/6J mice were intranasally infected with subclinical, non-encephalic HSV-1 (MaKrae strain) or mock infection, and brains were collected at 6 days post-infection (dpi), corresponding to peak innate immune activation^1^. Consistent with recent results, immunofluorescence revealed focal HSV-1 antigen localized predominantly to the spinal trigeminal nucleus (SpVN), consistent with viral entry and spread along trigeminal pathways (**Fig. 1B**) ^1, 29^. Consistent with our previous findings, antigen-positive regions showed increased Iba1 signal relative to mock and antigen-negative regions (**Fig. 1B–C**), and Iba1-positive area increased with HSV-1 signal (**Fig. 1D**). To quantify protein-level differences within these anatomically defined regions, we performed DSP-based spatial protein profiling using a 36-plex neural and glial antibody panel. Because viral antigen was confined to discrete ipsilateral regions, antigen-positive and antigen-negative ROIs were directly compared within the same infected sections compared to contralateral uninfected regions. ROI-level protein abundance differed between antigen-positive and antigen-negative regions (**Fig. 1E**). Normalized protein abundance values across the panel segregated by antigen status (**Fig. 1F**). ROI-wide protein signal also differed by antigen status (**S Fig. 1**). Proteins predominantly expressed in microglia and macrophage populations, including CD11b, CSTD, and MHC-II, exhibited higher ROI-level abundance in antigen-positive regions relative to antigen-negative regions (**Fig. 1G**). These markers reflect increased abundance of microglia actively involved in innate immune surveillance and antigen presentation. Their enrichment within antigen-positive ROIs is consistent with localized accumulation or activation of microglia and at spatially restricted sites of HSV-1 infection. This pattern suggests that HSV-1 infection elicits spatially restricted microglial activation, reflecting both proliferation of resident microglia and recruitment of peripheral macrophages to discrete regions of HSV-1 infection.

### Single-nuclei multi-omic profiling identifies transcriptionally and epigenetically distinct CD11b⁺ microglial populations

To define the cellular states underlying this localized microglial response, we performed single-nuclei multi-omic profiling of CD11b⁺ nuclei isolated from whole brains at 6 days post-infection. After quality control, paired transcriptomic (RNA) and chromatin (ATAC) accessibility profiles were obtained from 8,188 single nuclei (**Supplementary Fig. 2**). Unsupervised clustering of snRNAseq profiles identified eleven distinct cell populations. (**Fig. 2A**). These clusters were predominantly composed of CD11b⁺ monocytes (microglia and macrophages), alongside minor vascular and lymphoid populations (**Fig. 2A, left**). Marker gene-based annotation using established lineage markers distinguished multiple resident microglial activation states and infiltrating macrophage subsets, including ‘IFN-responsive’, ‘mitochondrial-activated’, and ‘immediate-early gene (IEG)-high microglia’ (**Fig. 2A, right**). Multiple transcriptionally distinct microglial states were resolved. The largest population comprised ‘homeostatic microglia’ (2,962 cells; 36% of total), defined by expression of canonical microglial identity markers including *P2ry12*, *Tmem119*, *Fcrls*, and *Gpr34*, which are widely used to define the homeostatic microglial state in mouse brain transcriptomic studies (**Fig. 1B; Supplementary Table 1A**) ^35,36^. Several activated microglia states were identified at substantial frequencies. ‘Transiently activated’ microglia represented 1,930 cells (23.6%) and were characterized by enrichment of immediate-early and regulatory genes *Jun*, *Btg2*, and *Zfp36*^37, 38^. IFN-responsive microglia comprised 894 cells (10.9%) and were defined by elevated expression of IFN-stimulated genes, including *Stat1, Stat2, Gbp2, Gbp5, Nlrc5, Ifi204,* and *Ifi207* (**Supplementary Table 1A**), with representative markers such as *Ccl12* and the immunity-related GTPase *Gm4951*, linking IFN signaling to inflammatory and metabolic pathways^39^. A population annotated as primed microglia (762 cells; 9.3%) was identified based on retained expression of broadly expressed microglial genes, including *Cst3, Hexb,* and *Tanc2*, in the absence of transcriptional programs characteristic of IFN-responsive, immediate-early, or metabolically activated microglial states. Mitochondrial-activated microglia (633 cells; 7.7%) were characterized by elevated expression of mitochondrial-encoded transcripts (*mt-Cytb, mt-Nd1, mt-Nd2*) alongside preserved microglial identity genes (*Cst3, C1qa, Csf1r*). Immediate- early gene (IEG)-high microglia were defined by strong induction of canonical activity-dependent transcription factors including *Egr1, Fos, Fosb, Jun,* and *Atf3*, along with rapid feedback regulators such as *Dusp1, Btg2,* and *Zfp36*.

**Figure 2.**
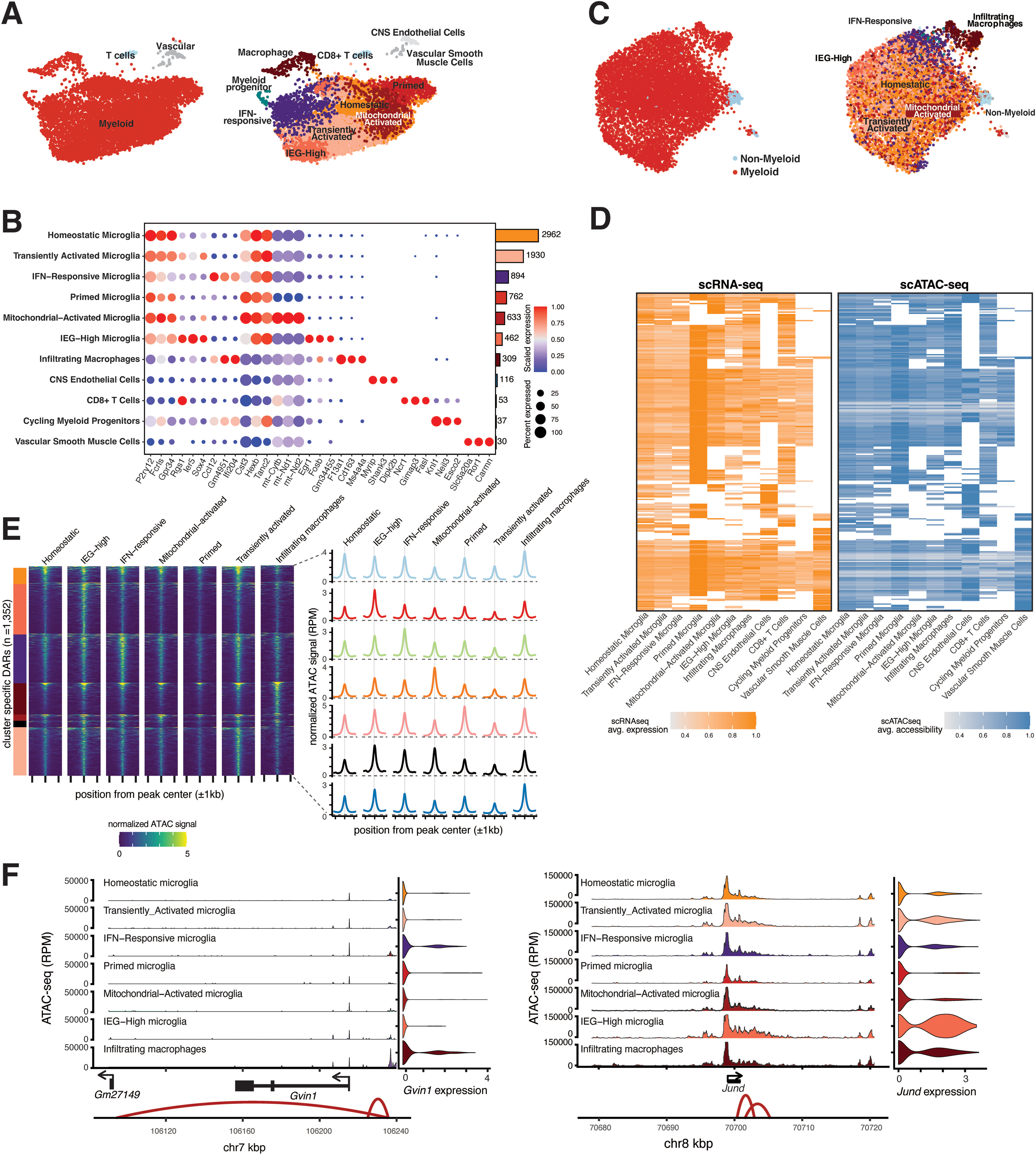
Single-nuclei multi-omic profiling reveals diverse microglial states and chromatin accessibility programs during acute HSV-1 infection. **A)** UMAP projection of CD11b⁺ nuclei based on the RNA modality (scRNA-seq), colored by broad (left; myeloid red non-myeloid light blue) and fine-grained (right) cluster annotations. **B)** Dot plot showing scaled expression (color) and percentage of expressing cells (dot size) for the top three marker genes per annotated cluster from the RNA modality. Bar plot (right) indicates the number of cells per cluster. **C)** UMAP projection of CD11b⁺ nuclei based on the ATAC modality (chromatin accessibility), colored by broad (myeloid red non-myeloid light blue) and fine-grained annotated clusters as in Fig. 1A. **D)** Heatmaps displaying matched single-cell RNA expression (left; light orange) and ATAC-derived gene activity scores (right; blue) for genes with the highest correlation between transcription and chromatin accessibility across clusters. Gene activity scores were inferred from scATAC-seq data by aggregating chromatin accessibility in promoter-proximal regions, spanning upstream of the transcription start site and across gene bodies. **E)** Heatmap showing normalized scATAC-seq signal across 1,352 cluster-enriched differentially accessible regions (DARs), centered at peak summits ±2 kb and grouped by annotated cluster. Right: average chromatin accessibility signal profiles for cluster-enriched DARs. Color bars correspond to annotated myeloid subtypes. **F)** Genome browser tracks showing normalized ATAC-seq signal (RPKM) across the *Gvin1* (chr7) and *JunD* (chr8) loci for each annotated cluster. Violin plots display log-normalized gene expression from the RNA modality for the same loci. Genomic coordinates are shown in kilobases (kb). Arcs represent predicted regulatory interactions between accessible chromatin regions and genes based on correlated accessibility patterns.

In addition to resident microglial states, a transcriptionally distinct population of infiltrating macrophages was identified, defined by high expression of canonical peripheral macrophage markers including *Mrc1, Cd163, Ms4a7, F13a1,* and *Dab2*, together with genes involved in cytoskeletal remodeling and migration (*Arhgap15, Vav3, Itga4*). A minor population of cycling CD11b^+^ cells was also detected (37 cells; 0.5% of total), characterized by strong enrichment of cell cycle and mitotic regulators, including *Mki67, Top2a, Knl1,* and *Cenpi* (**Supplementary Table 1A**).

We next examined chromatin accessibility in the same cells to assess whether the transcriptionally defined populations were also epigenetically distinct. Chromatin accessibility profiles largely mirrored the RNA-defined populations, with distinct separation of IFN-responsive microglia, infiltrating macrophages, and non-myeloid cells (**Fig. 1C**). In contrast, homeostatic, transiently activated, and mitochondrial-activated microglia showed more overlapping accessibility patterns. We identified 1,352 cluster-enriched differentially accessible regions (DARs) across annotated populations (**Fig. 1E**). Homeostatic and mitochondrial-activated microglia exhibited more discrete accessibility peaks, whereas IFN-responsive microglia and infiltrating macrophages displayed broader accessibility patterns. To relate chromatin accessibility to gene expression, we compared snATAC-derived gene activity scores with matched RNA expression across clusters (**Fig. 1D**). Many cluster-defining genes showed concordant accessibility and expression. For example, the IFN-induced gene *Gvin1* exhibited increased accessibility and RNA expression in IFN-responsive microglia (**Fig. 1F**), whereas the AP-1 transcription factor subunit *JunD* showed elevated accessibility and expression in IEG-high microglia.

### Primary HSV-1 infection induces transcriptional reprogramming and expansion of IFN-responsive microglia

To evaluate HSV-1 infection-associated changes in microglial populations, we compared single-nucleus RNA profiles from mock-treated and HSV-1-infected brains across annotated cell states. All major monocyte populations were detected in both conditions (**Fig. 3A**). However, their relative representation differed between mock and HSV-1 samples (**Fig. 3B**; **S Fig 3A**). IFN-responsive microglia were rare in mock-infected brains (11 nuclei; 0.47% of CD11b^+^ cells) but comprised 883 nuclei (15.7%) following HSV-1 infection. Infiltrating macrophages were detected in both conditions (116 nuclei, 4.97% in mock; 193 nuclei, 3.44% in HSV-1). Within the CD11b⁺ compartment, homeostatic microglia represented similar fractions across conditions (38.7% in mock; 36.6% in HSV-1). Transiently activated microglia were proportionally reduced in HSV-1 samples (33.8% vs 20.3%), whereas primed microglia were proportionally increased (5.7% vs 11.2%). Mitochondrial-activated and IEG-high microglia were present at comparable frequencies between conditions.

**Figure 3.**
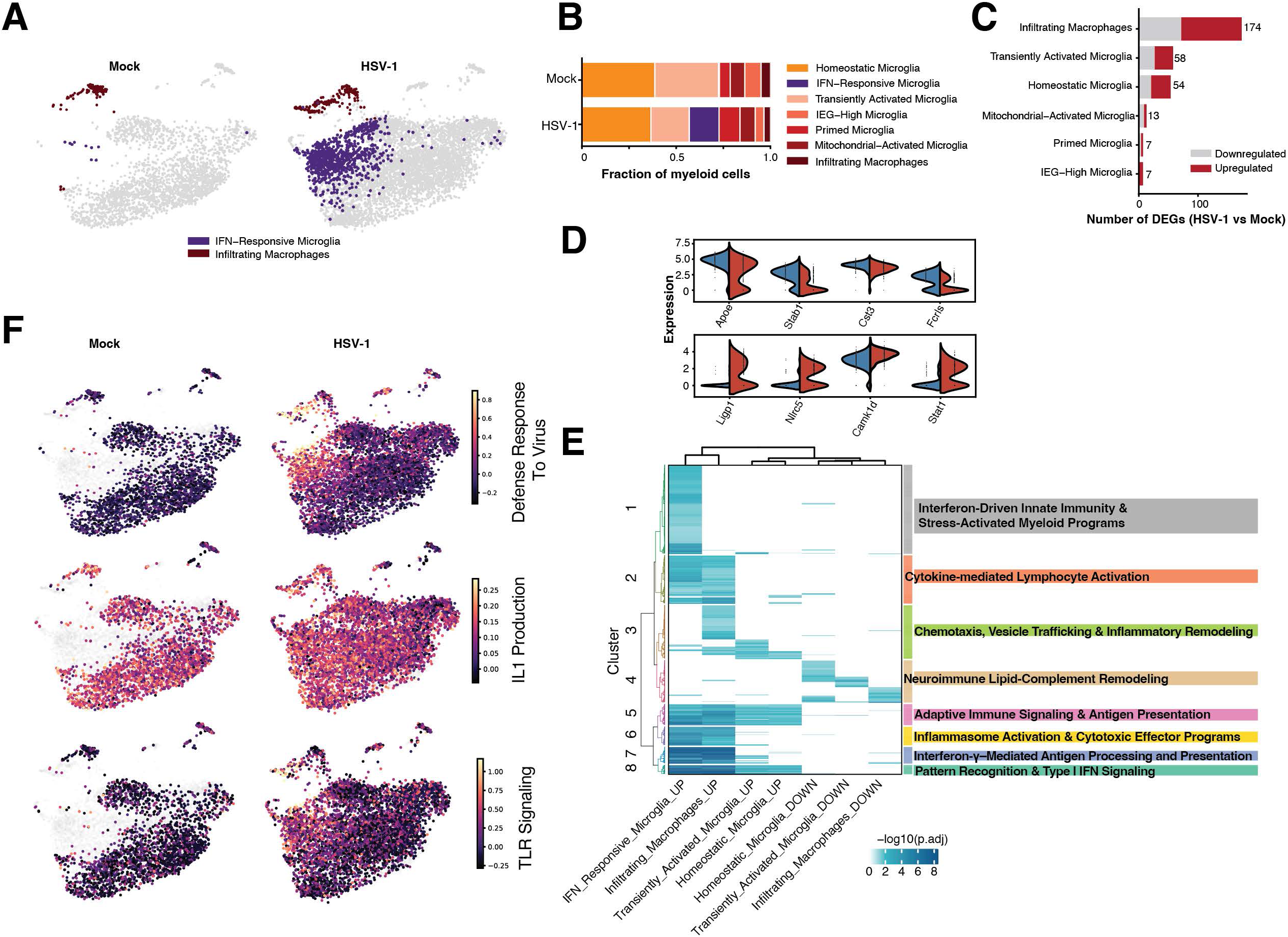
Primary HSV-1 infection remodels microglial states and regulatory programs in the CNS. **A)** UMAP of the RNA modality (snRNA-seq) separated by condition (Mock, HSV-1). IFN-responsive microglia and infiltrating macrophages are colored, and all other cells are shown in light gray. **B)** Relative abundance of annotated myeloid populations by condition. Stacked bars show the fraction of myeloid cells assigned to each annotated population within Mock and HSV-1 samples. **C)** Differential expression analysis across matched myeloid populations (HSV-1 vs Mock). The number of differentially expressed genes (DEGs) per annotated population is shown for populations with ≥50 nuclei per condition (adjusted p-value < 0.05 and |log2 fold-change| > 0.25). Red and gray bars indicate genes upregulated and downregulated in HSV-1, respectively. **D)** Representative DEGs in infiltrating macrophages comparing HSV-1 vs Mock split condition, Mock (blue) and HSV-1 (red). The y-axis shows log-normalized RNA expression values (normalized counts). Top and bottom rows indicate genes upregulated and downregulated in HSV-1, respectively. **E)** Over-representation analysis heatmap of enriched gene sets across differentially expressed gene lists. Columns correspond to DEG sets defined by annotated population and regulation direction (upregulated or downregulated genes), as well as the top 250 upregulated IFN-responsive microglia cluster marker genes. Rows are enriched gene sets (MSigDB C2, GO, KEGG, and Reactome). Heatmap color encodes enrichment significance as −log10(adjusted P value). Gene sets hierarchically clustered groups of similar gene sets that share similar enrichment patterns. 8 annotated pathway clusters (color strip at right). Cluster labels summarize the dominant biological programs represented within each group (Table S2). **F)** Pathway module scores projected onto RNA UMAP split by condition (Mock and HSV-1), for the gene sets Defense response to virus (GOBP), Interleukin-1 production (GOBP), Toll-like receptor signaling (KEGG). Module scores were computed using averaged expression of intersecting genes and color scales represent the module scores (low/high percentile limits).

To characterize HSV-1-associated gene expression changes across annotated populations, we performed differential expression analysis in matched microglial states with sufficient representation in both conditions, including infiltrating macrophages, transiently activated microglia, and homeostatic microglia (**Fig. 3C**). Infiltrating macrophages exhibited the largest number of differentially expressed genes (DEGs) following infection, with induction of antiviral regulators such as *Stat1*, *Nlrc5*, and *Ligp1*, and repression of homeostatic genes including *Cst3*, *ApoE*, *Stab1*, and *Fcrls* (**Fig. 3D**) ^40, 41^. To evaluate pathway-level responses, we performed over-representation analysis on stable microglial populations (≥50 cells per condition) exhibiting substantial gene expression changes (≥30 DEGs). Functional enrichment of HSV-1–associated DEGs, together with the top upregulated marker genes defining IFN-responsive microglia, identified eight pathway clusters spanning antiviral IFN signaling, inflammatory activation, antigen presentation, vesicle trafficking, and lipid–complement remodeling programs (**Fig. 3E**; **Table S2**) ^42^. Distinct MSigDB pathway patterns were evident across microglial states. Antiviral IFN-associated pathways (Clusters 1-2) were strongly enriched in both IFN-responsive microglia and HSV-1-associated infiltrating macrophages. The GOBP Defense response to virus gene set overlapped substantially with upregulated genes in infiltrating macrophages (19 genes) and IFN-responsive microglia (56 genes), including canonical IFN-stimulated genes (*Stat1*, *Stat2*, *Irf1*, *Irf7*, *Ifih1*, *Ifitm3*, *Eif2ak2*, *Samhd1*, *Zc3hav1*), chemokines (*Cxcl9*, *Cxcl10*), and guanylate-binding proteins (*Gbp2*, *Gbp5*, *Gbp7*) (**S Fig. 3B**) ^43–46^. Antiviral module scores were higher in IFN-responsive microglia and infiltrating macrophages in HSV-1 samples, with the strongest enrichment observed in the IFN-responsive microglial state (**Fig. 3G**). The GOBP Interleukin-1 production gene set overlapped predominantly with IFN-responsive microglia (14 genes), including inflammasome components and innate sensors (*Aim2*, *Nlrp3*, *Nod1*, *Tlr4*, *Il1a*) ^47–49^ and downstream mediators (*Jak2*, *Stat3*, *Smad3*), and showed minimal overlap in infiltrating macrophages (**Fig. 3G; S Fig. 3C**). The KEGG Toll-like receptor signaling pathway showed overlap with upregulated genes in infiltrating macrophages (*Akt3, Cxcl10, Cxcl9, Irf7, Stat1*). A similar set of TLR-associated genes was also detected in IFN-responsive microglia, including *Stat1* and *Stat2* (**S Fig. 3D**).

### Integration of RNA and chromatin accessibility identifies HSV-1–associated transcription factor regulons

To define regulatory programs associated with HSV-1 infection, we integrated RNA expression and chromatin accessibility using SCENIC+ to infer transcription factor regulons, defined as TFs and their predicted target genes supported by linked regulatory regions. Regulon activity in individual cells was quantified using AUCell enrichment scores together with motif accessibility. Across all annotated populations, regulon activity separated major cellular compartments, distinguishing microglial and macrophage cells (**S Fig. 4A**). Endothelial and vascular smooth muscle cells were enriched for ETS-family regulons (e.g., *Erg*, *Fli1*), whereas CD8⁺ T cells showed activity of lymphoid regulators such as *Lef1* and *Tbx21*.

**Figure 4.**
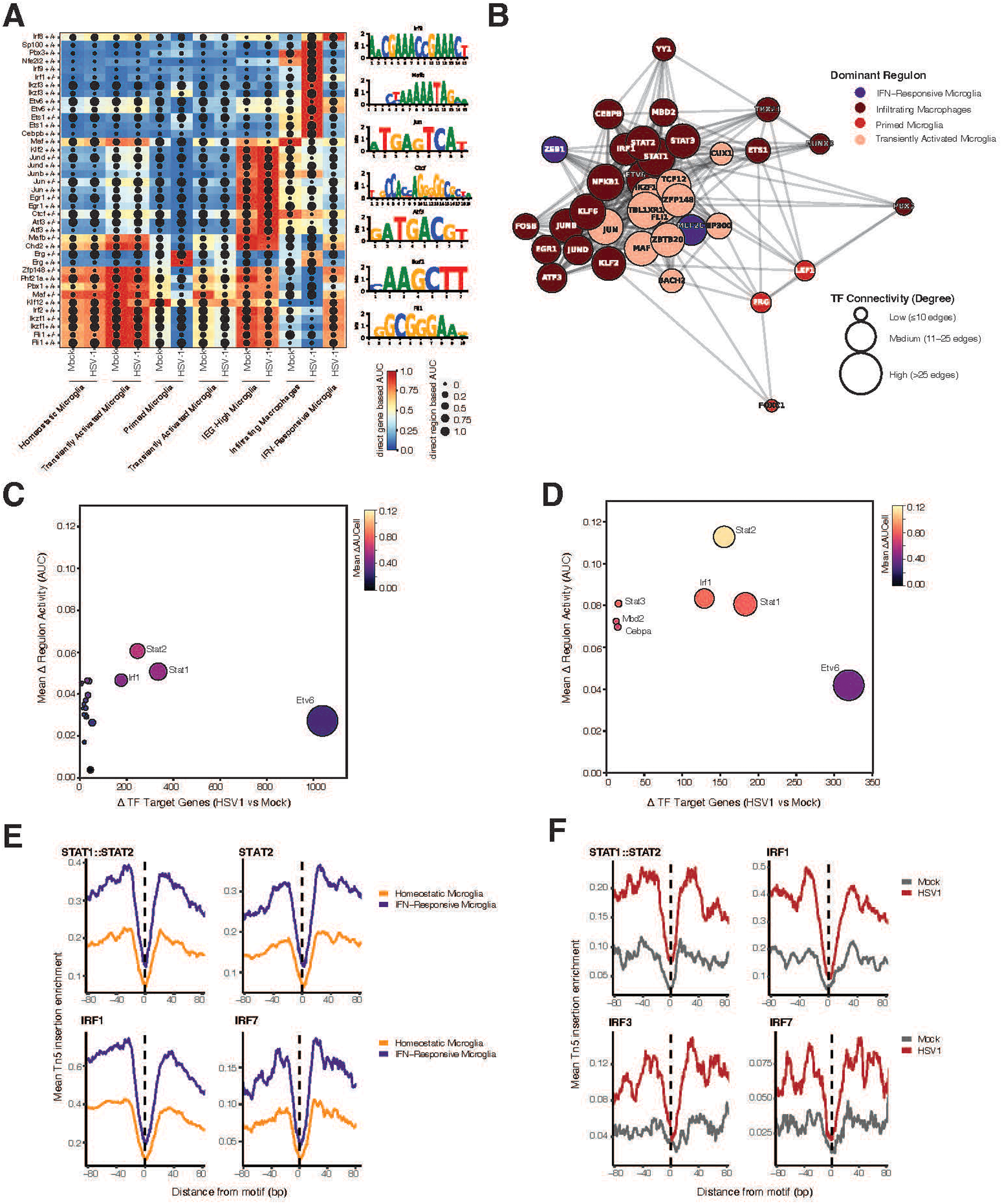
Transcription factor regulatory programs and connectivity in microglia and infiltrating macrophages in HSV-1-infected brains. **A)** Direct transcription factor regulon activity inferred using joint RNA+ATAC regulatory modeling with SCENIC+. Columns represent annotated myeloid populations split by condition (Mock, HSV-1). Rows represent direct TF regulons. Tile color indicates gene-based regulon activity (AUCell score). Dot size indicates region-based regulon activity derived from chromatin accessibility. Motif logos show representative DNA binding motifs corresponding to selected TF regulons. **B)** Transcription factor network summarizing inferred regulatory connectivity. Nodes represent TF regulons; edges represent shared inferred target genes. Node color indicates the myeloid state in which each TF shows the highest mean regulon activity. Node size reflects TF connectivity (degree), binned as indicated. **C)** Mean regulon activity shift (ΔAUCell; IFN-responsive minus homeostatic microglia) is plotted on the y-axis against the number of inferred target genes per transcription factor on the x-axis. Only TFs with significant activity shifts (padj < 1×10⁻¹⁰) are shown. Point size scales with the number of inferred target genes, and color encodes mean ΔAUCell. **D)** Parallel analysis in infiltrating macrophages comparing HSV-1 and mock conditions. Mean regulon activity shift (ΔAUCell; HSV-1 minus mock macrophages) is plotted against the number of inferred target genes per transcription factor. Point size reflects the number of inferred target genes, and color encodes mean ΔAUCell. **E)** Aggregate chromatin accessibility footprinting profiles for TF motifs in IFN-Responsive and Homeostatic microglial populations. Tn5 insertion enrichment is plotted across motif-centered windows (±80 bp) for differentially accessible regions (DARs) identified between IFN-responsive and homeostatic microglia (n=1275). Lines represent mean bias-corrected Tn5 insertion signal per group. The dashed vertical line marks the motif center (position 0). **F)** Footprinting profiles for TFs in infiltrating macrophages with HSV-1 infection. DARs identified between HSV-1 and mock conditions (n=883).

Within the microglial and macrophage compartment, transcription factor regulon activity differed across annotated states and between HSV-1 and mock conditions (**Fig. 4A**). IFN-responsive microglia showed strong induction of IFN-associated regulons, including *Irf1*, *Irf7*, *Stat1*, and *Stat2*. Infiltrating macrophages also showed elevated IFN regulon activity but were more prominently enriched for *Ap-1* family regulators (*Jun*, *Junb*, *Jund*, Atf3). IEG-high microglia displayed pronounced *Ap-1* activity, whereas homeostatic and primed microglia retained higher activity of lineage-associated regulators.

To examine regulatory connectivity, we constructed a transcription factor network based on shared regulon targets (**Fig. 4B**). IFN regulators clustered together, AP-1 factors formed a related but distinct module, and lineage-associated regulators showed broader connectivity across microglial states.

To identify regulators most strongly associated with the IFN-responsive microglial state, we compared IFN-responsive and homeostatic microglia and integrated regulon activity shifts with inferred target gene support (**Fig. 4C**). *Stat1*, *Stat2*, and *Irf1* exhibited both pronounced activity induction and extensive inferred target connectivity, distinguishing them from factors with smaller activity shifts or more limited target support. A parallel analysis in infiltrating macrophages comparing HSV-1 and mock conditions yielded similar induction of IFN regulators (**Fig. 4D**). In contrast, AP-1 factors showed substantial activity shifts but were associated with more moderate inferred target connectivity relative to STAT-family regulators. To determine whether inferred IFN regulons corresponded to increased transcription factor engagement at regulatory DNA, we quantified aggregate Tn5 insertion profiles centered on STAT and IRF motifs. Motif-centered depletion of insertions relative to flanking regions was used as a measure of transcription factor occupancy. In IFN-responsive microglia, STAT and IRF motifs exhibited increased footprint depth compared to homeostatic microglia, particularly at regions that gained accessibility in the antiviral state (**Fig. 4E**; **Fig. S4C**). Similarly, infiltrating macrophages showed enhanced STAT and IRF motif protection in HSV-1–infected samples relative to mock controls (**Fig. 4F**; **Fig. S4D**). Increased motif protection was most pronounced at regions that gained accessibility in the antiviral state, whereas regions that lost accessibility did not show comparable increases in footprint depth (Fig. S4A–B). These results indicate that chromatin remodeling at IFN-responsive regulatory elements is accompanied by increased engagement of STAT and IRF transcription factors, supporting the inferred regulatory circuitry.

### Spatial transcriptomics reveals localized IFN-driven immune activation and macrophage and microglial enrichment following HSV-1 infection

While single-nuclei multi-omic profiling defined state-specific induction of inflammatory transcription factor regulons within distinct macrophage and microglial populations, these analyses were performed on nuclei isolated from whole-brain tissue. However, our proteomic and histological analysis suggested these inflammatory signals may be spatially restricted. To determine whether the STAT- and IRF-associated gene programs identified at single-cell resolution were spatially restricted to sites of viral antigen, we analyzed transcriptomes from anatomically defined regions of interest (ROI). HSV-1 antigen–positive ROIs were confined to the spinal trigeminal nucleus (SpVN) and other spatially restricted regions of the brainstem in infected animals, but absent in mock controls. We focused on SpVN as the main route of HSV-1 neuroinvasion. Unsupervised projection of ROI transcriptomes showed segregation of HSV-1 antigen–positive regions from antigen-negative and mock ROIs (**Fig. 5A**). Differential expression analysis comparing HSV-1⁺ and HSV-1⁻ ROIs identified an induction of IFN-stimulated genes and inflammatory mediators (**Fig. 5B**). Among the most strongly upregulated genes were *Stat1*, *Ifit3*, *Irgm1/2*, *Oasl2*, and *Gbp3,* and activation of antiviral defense pathways. Hierarchical clustering of the top DEGs demonstrated coordinated induction of IFN-responsive gene modules in antigen-positive ROIs (**Fig. 5C**). Pathway enrichment analysis confirmed significant overrepresentation of IFN signaling, cytokine response, and viral sensing pathways among genes induced in HSV-1-positive regions (**Fig. 5D-E**), whereas there was a repression in metabolic and homeostatic pathways. To estimate infection-associated changes in cellular composition, we applied reference-based deconvolution of spatial transcriptomic profiles. HSV-1 antigen–positive ROIs showed reduced neuron- and astrocyte-associated expression signatures and increased microglial and macrophage signatures relative to antigen-negative regions (**Fig. 5F**). These spatial patterns are consistent with the expansion of IFN-responsive microglia and infiltrating macrophages defined by single-cell analysis. To directly test whether the regulatory architecture defined at single-cell resolution was spatially localized to infected tissue, we performed Fisher’s exact tests to assess enrichment of SCENIC-inferred transcription factor regulon targets among genes upregulated in HSV-1 antigen–positive ROIs (p < 0.05, |log2FC| > 0.5), using all expressed genes as the background universe. Genes induced in HSV⁺ ROIs were strongly enriched for targets of STAT2 (OR = 12.2, FDR = 9.6 × 10⁻²⁹), IRF1 (OR = 10.7, FDR = 2.7 × 10⁻²³), and STAT1 (OR = 7.4, FDR = 1.5 × 10⁻¹⁹) regulons, with similarly elevated enrichment for STAT3 and CEBPB targets (**Fig. 5G**). In contrast, these HSV-1–associated regulons were not enriched among genes downregulated in HSV⁺ ROIs (OR ≤ 0.13, FDR > 0.05), indicating that spatial transcriptional activation reflects selective induction of the same regulatory programs identified in HSV-responsive microglial states.

**Figure 5.**
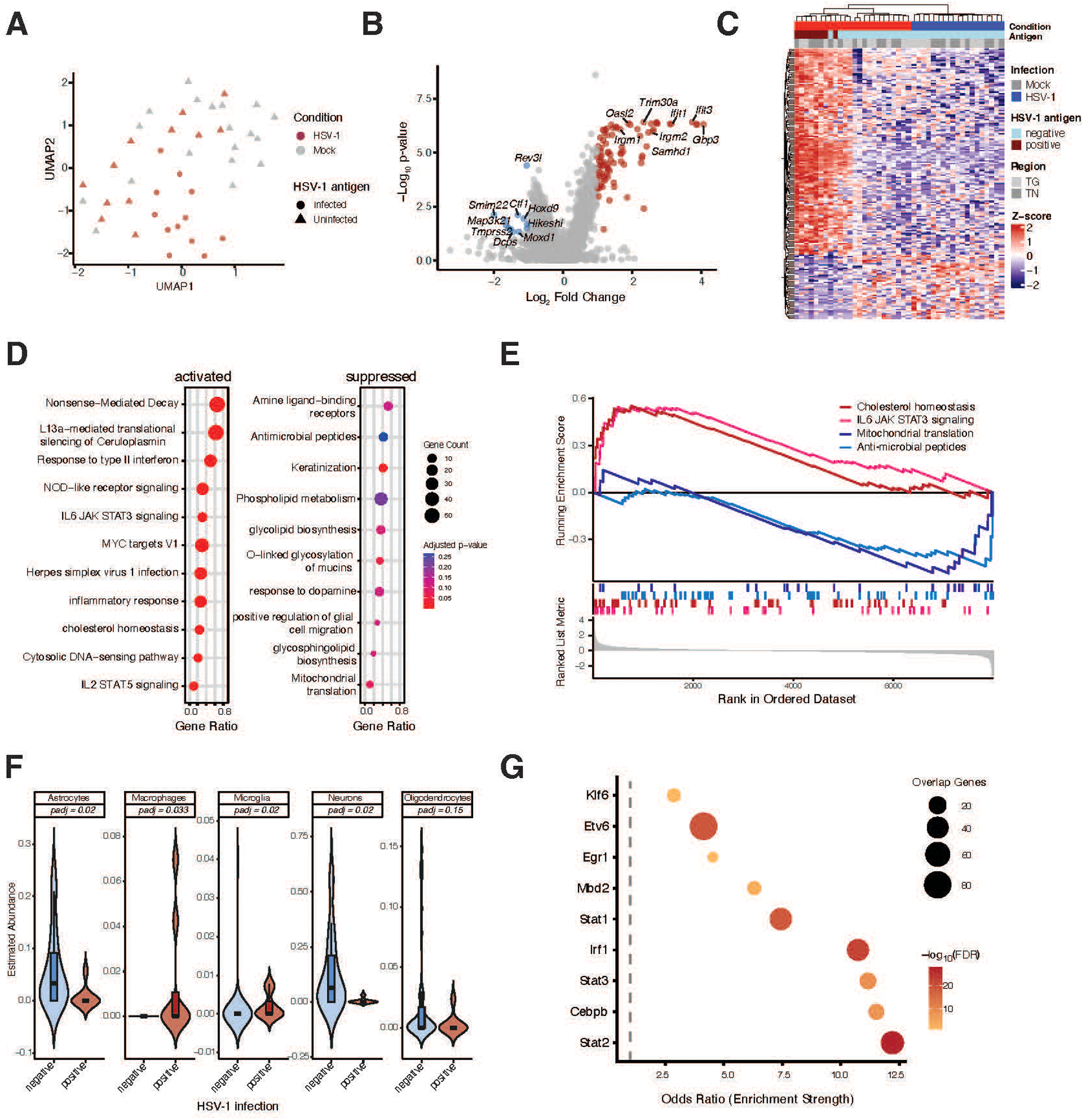
Spatial transcriptomic profiling identifies localized transcriptional activation and enrichment of HSV-1–associated transcription factor regulon targets in HSV⁺ brainstem regions. **A)** UMAP projection of spatial whole-transcriptome profiles from all regions of interest (ROIs). Points represent individual ROIs and are colored by infection condition (HSV-1, red; mock, gray). Shape denotes HSV-1 antigen status (circles, antigen-positive; triangles, antigen-negative). **B)** Differentially expressed genes (DEGs) between HSV-1 antigen-positive and antigen-negative ROIs. Red and blue points represent significantly upregulated and downregulated genes, respectively (log₂FC > 0.5, p < 0.05). Top DEGs are labeled. **C)** Heatmap of the top 100 differentially expressed genes across all ROIs. Columns represent individual ROIs and rows represent genes. Expression values are Z-score scaled per gene. Annotation bars indicate infection condition (HSV-1 or mock), HSV-1 antigen status (positive or negative), and anatomical region (trigeminal ganglion, TG; trigeminal nucleus, TN). **D)** Pathway enrichment analysis of genes differentially expressed in HSV-1 antigen–positive ROIs. Left panel shows pathways enriched among upregulated genes; right panel shows pathways enriched among downregulated genes. Dot size indicates gene ratio (proportion of genes within the pathway represented in the DEG list), and color indicates adjusted p-value. **E)** Gene set enrichment analysis (GSEA) plots for representative pathways. Running enrichment scores are shown for IL6/JAK/STAT3 signaling, cholesterol homeostasis, mitochondrial translation, and antimicrobial peptides. The x-axis represents gene rank in the ordered dataset and the y-axis shows the running enrichment score. **F)** Cell type abundance estimates derived from DSP expression data using deconvolution with the mouse cell atlas reference, showing the estimated abundance of major CNS cell types across HSV-1 antigen-positive and antigen-negative ROIs. Differences were assessed using Wilcoxon rank-sum tests; adjusted p-values (FDR) are shown. **G)** Enrichment of SCENIC-inferred transcription factor regulon target genes among genes upregulated in HSV-1 antigen–positive ROIs. Each point represents a transcription factor. The x-axis shows odds ratio from Fisher’s exact test. Point size reflects the number of overlapping genes between the regulon target set and spatially upregulated genes. Point color indicates −log₁₀(FDR). Only transcription factors with FDR < 0.05 are shown.

## Discussion

In this study, we define the spatial and regulatory architecture of monocyte and microglia responses during acute HSV-1 infection. By integrating spatial transcriptomics with single-nucleus RNA and chromatin accessibility profiling, we show that intranasal HSV-1 infection induces anatomically defined innate immune activation within discrete brainstem regions. Regions containing viral antigen exhibit strong induction of IFN-stimulated genes, enrichment of microglial and macrophage transcriptional signatures, and reduced neuronal and astrocytic transcript abundance. At single-cell resolution, we identify distinct microglial states, including an expanded IFN-responsive population characterized by increased STAT1/2- and IRF-centered regulon activity and enhanced chromatin accessibility at IFN-responsive regulatory elements. Together, these data show that acute HSV-1 infection elicits spatially confined, state-specific activation of microglial and macrophage populations rather than uniform CNS-wide inflammation. Spatially resolved protein profiling reinforced this conclusion. Within infected brains, direct comparison of antigen-positive and antigen-negative regions from the same tissue sections demonstrated increased abundance of microglial activation markers (CD11b, CTSD, MHC II) specifically in antigen-positive regions. These protein-level changes confirm that localized antiviral transcriptional activation is accompanied by measurable increases in microglial protein abundance and activation at sites containing viral antigen.

Our findings extend prior observations that HSV-1 infection produces focal microglial activation within the brainstem^1^. Viral antigen was confined to discrete regions of the ipsilateral spinal trigeminal nucleus, and these antigen-positive areas exhibited elevated Iba1 signal and strong correlation between viral burden and microglial activation. Spatial transcriptomic profiling revealed that HSV-1 antigen–positive regions are transcriptionally distinct from antigen-negative regions within the same infected tissue sections, with marked induction of canonical IFN- stimulated genes, including *Stat1, Irf7, Ifit3*, and *Gbp*. Importantly, these transcriptional changes were anatomically constrained. Rather than a diffuse inflammatory response, antiviral gene induction was localized to regions containing viral antigen. Deconvolution analysis further indicated enrichment of microglial and macrophage gene signatures in antigen-positive regions, with relative reduction of neuronal and astrocytic transcripts. Although reduced neuronal and astrocytic signatures may reflect altered gene expression rather than definitive cell loss, these findings indicate that HSV-1 neuroinvasion perturbs local cellular composition and transcriptional balance in a regionally restricted manner. This shift in regional cellular composition is consistent with previous reports that neurons and astrocytes are vulnerable to HSV-1 infection, and the reduction of neuronal and astrocytic transcriptional signatures in infected regions may reflect early degeneration or cellular stress in response to viral invasion^50,51^. Such localized disruption of neuroglial homeostasis supports the concept that HSV-1 infection drives focal immune remodeling that may predispose specific brain regions to long-term neuropathology.

Single-nucleus RNA and ATAC profiling identified multiple transcriptionally distinct microglial populations. Among these, an IFN-responsive microglial state was markedly expanded following infection, consistent with earlier studies^29, 52^. This population was characterized by elevated expression of *Stat1, Stat2, Irf1, Irf7*, and multiple IFN-stimulated and antiviral genes. The prominence of STAT1/2-centered programs is consistent with the established role of type I IFN signaling in antiviral defense^46, 53, 54^. In support, STAT1 deficiency is associated with increased susceptibility to HSV-1 infection in both murine and human systems, underscoring the importance of this pathway in host protection^44, 55–58^. Given that IFN-responsive microglial populations were rare in mock-infected brains, and predominantly associated with HSV-1-infected brains, it is reasonable to suggest that elevated IFN -dominant states, if unchecked, may eventually predispose the brain to neuroinflammatory and neurodegenerative disease^34^.

Our findings reveal that IFN-dominant states are epigenetically regulated and potentially sustained, with STAT- and IRF-centered regulatory programs selectively amplified during acute CNS infection. Specifically, we show that chromatin accessibility is increased at regulatory regions enriched for STAT and IRF motifs, and motif footprinting demonstrated enhanced transcription factor engagement at these sites. These findings clearly demonstrate that HSV-1 infection drives the epigenetic remodeling of IFN-dominant microglial states. Given that elevated IFN signaling promotes microglial dysfunction and is consistently linked to neurodegenerative pathology^59^, epigenetic maintenance of IFN-dominant states may underlie the persistent microglial dysfunction that precedes neurodegenerative disease.

Infiltrating macrophages also exhibited IFN-associated regulon activity, although with additional enrichment of *Ap-1*–associated programs, suggesting that resident and recruited microglial and macrophage populations deploy overlapping but non-identical regulatory circuits. The prominence of *Ap-1* is also notable given its emerging role as a functionally dynamic regulator of glial responses to CNS injury and degeneration^60^. In other contexts, transient AP-1 activation appears to support protective glial functions, including inflammatory coordination, debris clearance, and tissue repair, whereas persistent *Ap-1* activity has been linked to chronic microglial activation, synapse remodeling, tissue loss, and neurodegenerative pathology^61^. In this framework, AP-1 activity in infiltrating macrophages may reflect an acute host-defense response to HSV-1 neuroinvasion, which, if sustained, could contribute to prolonged immune activation and tissue vulnerability.

A key strength of this study is the integration of single-cell regulatory modeling with spatial transcriptomics and proteomics. Genes upregulated in HSV-1 antigen–positive regions were strongly enriched for predicted targets of *Stat1-, Stat2-, Irf1-, and Cebpb*-associated regulons identified at single-cell resolution. This spatial enrichment indicates that regulatory programs inferred from isolated nuclei correspond to transcriptional activation within anatomically defined regions *in vivo*. The accompanying increase in microglial activation and astrocytic proteins within antigen-positive regions further supports the conclusion that IFN-driven regulatory programs translate into tissue-level changes. While single-cell profiling defines the regulatory architecture of microglial states, spatial approaches confirm that these programs are geographically restricted to regions containing viral antigen.

Although IFN-responsive microglia were markedly expanded following infection, a homeostatic microglial population remained detectable. Several canonical homeostatic genes, including *Cst3, Stab1,* and *Fcrl*s were reduced in IFN-dominant states relative to the homeostatic cluster, suggesting a redistribution of microglial identities rather than the complete elimination of homeostatic microglia. Nonetheless, the transition away from homeostatic microglial states may have important functional consequences. For example, *Stab1* and *Fcrls* encode scavenger receptors with key roles in receptor-mediated endocytosis and recycling^62, 63^. *Cst3* is thought to inhibit cysteine proteases, induce autophagy, and inhibit Aβ oligomerization and fibrilization, and is therefore neuroprotective, especially in the context of AD^64^ ^65^. Collectively, the suppression of these factors could impair microglial phagocytosis, clearance of neurotoxic factors, and promote protein aggregation, contributing to increased risk of neurodegenerative disease following HSV-1 infection.

Our data also demonstrate coordinated suppression of homeostatic microglial signatures, characterized by *P2ry12*, *Tmem119*, and *Cx3cr1*, which are critical for microglial surveillance, synaptic remodeling, and neuronal communication^66^ ^67–69^. Moreover, viral infection and antiviral programs actively repress genes associated with lipid metabolism (*Trem2, ApoE,* and *Tyrobp*) ^70,71^ ^72^, transcripts which are often elevated in an attempt to achieve homeostasis in the aged and diseased brain. Downregulation of these pathways suggests that HSV-1 infection shifts microglia away from their homeostatic maintenance functions toward a highly specialized antiviral state. Loss of these homeostatic programs has been reported in multiple neurological disorders, including Alzheimer’s disease and multiple sclerosis ^73–75^, raising the possibility that viral infection may initiate microglial state transitions that resemble those observed in neurodegenerative conditions.

Chromatin accessibility profiling revealed that IFN-associated transcriptional activation is accompanied by increased accessibility at regulatory regions containing STAT and IRF motifs. Regions that gained accessibility in IFN-responsive microglia exhibited enhanced motif footprinting, consistent with transcription factor occupancy. These data indicate that antiviral gene induction is supported by coordinated changes in chromatin accessibility consistent with recent studies showing sustained IFN signaling and prolonged STAT1 activation initiate chromatin remodeling at antiviral loci in microglia^70, 76^. While our analysis does not directly measure histone modifications, the concordance between increased accessibility, regulon activity, and transcriptional induction suggests that acute HSV-1 infection engages a defined antiviral regulatory architecture in microglia. Whether these accessibility changes persist beyond the acute phase remains an important question for future study. Since HSV-1 is well-known to reactivate within the CNS^77^, exacerbating disease risk^3, 78^, it is plausible to speculate that initial HSV-1 infection increases chromatin accessibility at IFN-regulated loci, epigenetically maintaining immune memory, which amplifies the antiviral or inflammatory responses upon subsequent HSV-1 challenge.

Microglia are generally resistant to productive HSV-1 infection *in vivo*, and our data are consistent with a model in which many microglia mount IFN-driven responses without evidence of widespread viral gene expression or cell death signatures^79^. The presence of strong IFN signaling alongside limited viral transcript detection supports the possibility of bystander or abortive activation. However, this observation may reflect a selection bias resulting from the acute loss of vulnerable microglia, an effect that cannot be definitively resolved in the current dataset. However, future studies that definitively determine microglial infection status *in vivo* are warranted and will require targeted approaches to detect viral genomes and transcripts at single-cell resolution.

Our analysis was conducted at a single acute time point (6 days post-infection), capturing peak innate immune activation. This design limits assessment of whether the IFN-responsive states and associated chromatin accessibility changes resolve, persist, or evolve during later stages of infection, latency, or reactivation. Longitudinal profiling will be necessary to determine the durability of these microglial and macrophage state transitions. Additionally, our spatial analyses focused on the brainstem region corresponding to trigeminal entry into the CNS. However, our previous findings suggest multiple vulnerable regions of the brain during primary infection and a heterogeneous microglial response^1, 12, 80–82^. Whether similar regulatory programs are engaged in other brain regions during broader viral spread or reactivation remains to be determined. Expanding the anatomical scope of analysis will clarify whether the antiviral regulatory architecture we describe represents a generalizable CNS response or a regionally specialized phenomenon. Determining the impact of HSV-1 infection on microglial function is also warranted. While transcriptional and accessibility data indicate redistribution of microglial states and induction of antiviral programs, the consequences for phagocytosis, synaptic remodeling, and cytokine production remain to be directly tested.

Given the growing evidence linking HSV-1 to long-term neuroinflammatory phenotypes and Alzheimer’s disease pathology, it is critical to explore the durability of these microglial state transitions. Longitudinal spatial and single-cell profiling, combined with neuropathological assessments, will be essential to understand whether the observed loss of homeostatic signatures and amplification and maintenance of antiviral states contribute to persistent neuroinflammation and increased risk of neuroinflammatory and neurodegenerative disease.

Importantly, our findings provide a potential mechanistic framework linking acute viral infection to later-life neurological disease. If IFN-driven microglial states become epigenetically stabilized following viral exposure, repeated viral reactivation events could reinforce these inflammatory programs over time. Assessing how repeated reactivation episodes affect these microglial trajectories remains a key unanswered question. Such cumulative immune remodeling may contribute to regionally selective neurodegeneration, particularly in brain regions repeatedly exposed to viral entry or reactivation events. Additionally, these spatially defined immune signatures may also expand during neurodegenerative diseases. Therefore, future students should determine how viral infections reshape microglial regulatory states across disease contexts to elucidate how latent neurotropic viruses contribute to chronic neuroinflammation and heightened susceptibility to neurodegenerative disease.

## Conclusion

In summary, this study integrated a spatial and single-cell framework for understanding microglial responses to acute CNS HSV-1 infection. We demonstrate that antiviral IFN programs are induced in anatomically confined brainstem regions containing viral antigen and are associated with expansion of IFN-responsive microglial states characterized by distinct transcription factor regulon activity and chromatin accessibility patterns. These findings define the spatial and regulatory organization of early HSV-1–associated neuroinflammation and establish a foundation for investigating how recurrent viral exposure may shape long-term CNS immune states and disease susceptibility.

## METHODS

### Animals

Adult C57BL/6J mice (Jackson Laboratories) were housed under a 14-hour light/10-hour dark cycle with ad libitum access to food and water. All procedures were approved by the Institutional Animal Care and Use Committee at the University of Colorado Anschutz Medical Campus and conducted in accordance with NIH guidelines.

### In Vivo Model of HSV-1 Infection

Intranasal HSV-1 infection was performed as previously described^1, 83^. Briefly, mice were anesthetized with isoflurane and inoculated intranasally with 20 μL HSV-1 McKrae strain (1×10⁶ plaque-forming units per animal; GenBank accession JX142173). Mock-infected controls received vehicle alone. Animals were sacrificed at 6–7 days post-infection (dpi). Brainstem tissue was harvested for spatial transcriptomics or CD11b+ microglial isolation for single-nucleus multiome analysis.

### Spatial Transcriptomics (GeoMx DSP) and Proteomics (nCounter)

Mice were deeply anesthetized and transcardially perfused with PBS followed by 4% paraformaldehyde (PFA). Brainstems were post-fixed in 4% PFA (24–48 hours, 4°C), cryoprotected in 30% sucrose, embedded in OCT, sectioned at 5–10 μm, and stored at −80°C. Sections were processed for immunofluorescence to identify regions of interest (ROIs). Antigen retrieval was performed using heated citrate buffer, followed by permeabilization in 1% Triton X-100 (1 hour), blocking in 5–10% normal serum (goat or donkey) with 0.1% Triton X-100 (1 hour), and overnight incubation with primary antibodies in PBS containing 0.1% Triton X-100 and 1% serum. Primary antibodies included rabbit anti–HSV-1 glycoproteins (DAKO, B011402-2, 1:1000) and goat anti-Iba1 (Abcam, ab5076, 1:500). Secondary antibodies were applied for 2 hours followed by DAPI counterstaining.

Selected sections were processed using the NanoString GeoMx Digital Spatial Profiler (DSP) Whole Transcriptome Atlas (WTA) assay with SYTO13 nuclear stain and HSV-1/Iba1 immunofluorescent labeling. ROIs were manually selected based on anatomical location and HSV-1 antigen intensity. Antigen-positive and antigen-negative regions were analyzed from the same sections when applicable. Raw WTA counts were processed using GeoMxTools (NanoString). Data were quantile normalized (q_norm) and log₂-transformed (log_q). ROIs with low nuclei counts or low gene detection were excluded. Differential expression was performed using Wilcoxon rank-sum tests on log_q values, retaining genes with |log₂ fold change| > 1 and p < 0.05. Gene set enrichment analysis (GSEA) was performed using genekitr (v1.0.4) against KEGG, Reactome, and Hallmark gene sets, with FDR < 0.025 considered significant. Cell type deconvolution was performed using spatialdecon (v1.10.0) with the Mouse Cell Atlas adult brain reference matrix. Cell types were collapsed into six categories (astrocytes, microglia, neurons, oligodendrocytes, endothelial cells, pericytes) ^84^. Statistical comparisons were performed using Wilcoxon or Kruskal–Wallis tests in R (v4.3.2).

Spatial protein expression profiling was performed using the NanoString GeoMx Digital Spatial Profiler (DSP), utilizing the Mouse Glial Cell Subtyping Protein Module 1.0 and Mouse Neural Cell Profiling Protein Core 1.0 panels. Protein counts were processed using the standard GeoMx DSP normalization pipeline, and all ROIs passed QC. Normalized counts were log2-transformed (log2[counts + 1]) prior to downstream analysis. Differential protein abundance between HSV-1 antigen–positive and antigen–negative regions of interest (ROIs) was assessed using linear modeling with the limma framework, including antigen status as the sole variable. Proteins with adjusted p-value < 0.05 were deemed significant. To evaluate the contribution of global shifts in protein abundance, a secondary analysis was performed using per-ROI median-centered expression values. No additional background correction was applied beyond the standard GeoMx DSP normalization workflow.

### Microglia Isolation

Mice were euthanized, and tissues were perfused with HBSS according to institutional protocols. Brains were excised and finely minced in digestion medium (Papain, DNase 1, Earls Balanced Salt Solution [EBBS]). A single-cell suspension was achieved by digesting at 37 °C for 30 minutes with titrations every 5-10 minutes, reducing the size of the pipette each time (e.g., 10 ml, 5 ml, 1 ml). Samples were then diluted to prevent further digestion and filtered through a 70μM cell strainer. The flow-through was then centrifuged at 300g for 10 minutes at 4 °C with minimal braking. The supernatant was discarded, and the cell pellet was resuspended in 1 ml HBSS (2% FBS). To isolate microglia from the mixed cell suspension, cells were incubated with CD11b+ magnetic beads according to the manufacturer’s instructions (Stem cell, Easy Sep), and incubated with a magnet. CD11b- cells were removed by tipping the magnet and supernatant. CD11b+ cells were washed three times before being eluted off the magnet.

### Isolation of Nuclei

The number of cells in the CD11b+ enriched microglia fractions was quantified, cells were pelleted (500g for 5 mins) and resuspended at 100,000 per 100 μl. Cells were allowed to lyse on ice for 5 minutes in cell lysis buffer (10 mM Tris-HCl pH7.4, 10 mM Nacl, 3 mM MgCl2, 0.1% Tween-20, 0.01% IGEPAL CA-630, 0.01% Digitonin, 1% BSA, 1% DTT, 1 U/ML RNase inhibitor). 1 ml of chilled wash buffer (10 mM Tris-HCl pH7.4, 10 mM Nacl, 3 mM MgCl2, 0.1% Tween-20, 1% BSA, 1% DTT, 1 U/ML RNase inhibitor) was added to the cells. To remove unwanted debris and magnetic beads, the lysed cells were incubated with the magnet for 3 minutes on ice. The supernatant, containing the nuclei, was then poured off, while the cell debris remained attached to the magnet. The nuclei were then pelleted at 500g for 5 mins and washed three times in nuclear buffer (1x Nuclear buffer [Multiome ATAC + GEC kit] 1 mM DTT, 1 U/ml RNase inhibitor). After the final wash, cells were resuspended in at least 15 μl of nuclear buffer at a concentration of 1200 nuclei/μl ready, the optimal concentrations for multiome snRNAseq and snATACseq experiments.

### Single-Nucleus Multiome Library Preparation and Sequencing

Single-nuclei RNA and ATAC libraries were prepared using the 10x Genomics Chromium Single-nuclei Multiome ATAC + Gene Expression kit following the manufacturer’s protocol. Libraries were sequenced on an Illumina platform to achieve recommended depth for both gene expression and chromatin accessibility. Reads were aligned to the mm10 reference genome using Cell Ranger ARC (10x Genomics). RNA and ATAC fragments were jointly processed to generate count matrices and fragment files for downstream analysis.

### Single-Nucleus RNA-seq Analysis

RNA data were analyzed using Seurat (v5). Cells were filtered based on gene number, UMI counts, and mitochondrial percentage. Specifically, cells with low gene detection (<200 genes), extreme gene counts, or >10% mitochondrial transcripts were excluded. Data were normalized, variable features identified, scaled (regressing percent.mt), and subjected to PCA. Nearest-neighbor graphs were constructed using the first 10 principal components followed by clustering (resolution 0.4) and UMAP visualization. Cluster identities were assigned based on canonical marker genes and module scoring. Differential expression between HSV-1 and mock conditions was performed using Wilcoxon rank-sum testing with adjusted p < 0.05 and log₂ fold change > 0.25.

### Single-Nucleus ATAC-seq Analysis

ATAC fragments were analyzed using Signac and pycisTopic (v2.0a0). Cells were filtered based on unique fragment counts and transcription start site (TSS) enrichment scores. Fragment size distributions and TSS enrichment profiles were examined to confirm data quality. Pseudobulk accessibility profiles were generated per cell type and condition using export_pseudobulk. Peaks were called using MACS2 (v2.2.9.1) with no model building, 73 bp shift, 146 bp extension, duplicates retained, and q < 0.05. NarrowPeak outputs were merged into a nonredundant consensus peak set after removing ENCODE blacklist regions. A sparse cell-by-region matrix was constructed and modeled using Latent Dirichlet Allocation (LDA) with Mallet. Models spanning 2–50 topics were evaluated; a 45-topic model was selected. Differentially accessible regions (DARs) were identified using find_diff_features (adjusted p < 0.05, log₂FC > log₂1.5). Gene activity scores were calculated by linking accessible regions within gene bodies or ±100 kb of transcription start sites.

### SCENIC+ eGRN Inference

SCENIC+ (v1.0a2) was used to integrate chromatin accessibility and gene expression data for eRegulon inference. Candidate regulatory regions were defined within ±150 kb of TSSs (excluding the first 1 kb). Motif enrichment was performed using mm10 SCREEN v10 motif databases. Significant motifs were retained using FDR < 0.001, NES ≥ 3.0, and AUC ≥ 0.005 thresholds. Gradient boosting models and Spearman correlation were used to score TF–region and region–gene relationships. eRegulons were retained if ≥10 target genes were identified with correlation ρ > 0.05. AUCell was used to quantify per-cell regulon activity. Target genes of *Irf1, Stat1, Stat2, Stat3, and Mbd2* were further analyzed to generate TF–region–gene network visualizations using NetworkX. Module scores were computed using Scanpy’s score genes function on matched RNA data.

### Statistics

Spatial transcriptomics, multiome, and pathway analyses are described above. All additional statistical analyses were conducted using GraphPad Prism. Comparisons between two groups were performed using either parametric or nonparametric *t*-tests, as appropriate. Comparisons involving more than two groups were analyzed using one-way analysis of variance (ANOVA). Exact *P* values are reported, with statistical significance defined as *P* < 0.05.

### Study protocols

All animal procedures were approved by the Institutional Animal Care and Use and Ethics Committee at the University of Colorado Anschutz Medical Campus and conducted in accordance with NIH guidelines.

## Acknowledgements

We would like to thank the Genomics Shared Resources Core (P30CA046934) at the University of Colorado, Anschutz Medical Campus, for their technical support. We would also like to thank the personnel from the University of Colorado Anschutz Medical Campus Advanced Light Microscopy Core for imaging assistance. Imaging was performed in the Advanced Light Microscopy Core facility of the NeuroTechnology Center at the University of Colorado Anschutz Medical Campus, which is supported in part by Rocky Mountain Neurological Disorders Core Grant (P30NS048154) and by Diabetes Research Center Grant (P30 DK116073).

## Disclosures

The authors have nothing to declare

## Study Funding

This research was supported by grants from NIH/NCATS Colorado CTSA K12 TR004412 (C.S.N.), an Across the Finish Line (AFL) grant from the University of Colorado School of Medicine awarded to K.D.B and C.S.N, an NIH NIA R01 AG079217 (K.D.B and D.O), an NIH NIA R21 awarded to K.D.B (R21AG091650), NIH NIA R01 AG079193 (D.R.; M.A.N; C.S.N, A.N.B, A.B.S.), NIH NIAID R21 AI176110 (A.N.B).

## Author Contributions (CRediT taxonomy)

**S.F.**: Conceptualization; Formal analysis; Data curation; Writing – original draft; Writing – review & editing.

**C.L.**: Formal analysis; Data curation; Writing – review & editing.

**D.O.**: Investigation.

**J.B.**: Formal analysis.

**A.N.B.**: Data interpretation; Writing – review & editing.

**A.S.B.**: Investigation.

**M.A.N**: Data interpretation; Writing – review & editing.

**D.R.**: Data interpretation; Writing – review & editing.

**K.D.B.**: Conceptualization; Methodology; Investigation; Writing – original draft; Writing – review & editing.

**C.S.N.**: Conceptualization; Methodology; Investigation; Writing – original draft; Writing – review & editing.

## Data availability

Raw sequencing reads are available through the Sequence Read Archive and are linked to the corresponding GEO records, and processed count matrices are provided within the GEO submissions. GSE324176 NanoString GeoMx Whole Transcriptome Atlas spatial transcriptomics dataset. GSE324144 Single-cell multiome (scRNA-seq + scATAC-seq) dataset from HSV-1 infected and mock mouse brain.

---

**Supplementary Figure 2. Quality control and filtering of single-nuclei multiome data.**

**A)** Barcode rank (knee) plots for mock and HSV-1–infected samples showing total UMI counts per barcode. Dashed lines indicate the selected inflection threshold used to define high-quality nuclei retained for downstream analysis. The number of retained nuclei per condition is indicated.

**B)** Violin plots showing distributions of detected genes (nFeature_RNA), total RNA counts (nCount_RNA), and mitochondrial transcript percentage (percent.mt) across mock and HSV-1 samples prior to filtering.

**C)** Scatter plots of RNA quality-control metrics. Left: nFeature_RNA versus nCount_RNA illustrating the relationship between library complexity and sequencing depth. Middle: percent.mt versus nCount_RNA. Right: percent.mt versus nFeature_RNA. Dashed lines indicate applied filtering thresholds for gene detection and mitochondrial transcript content.

**D)** ATAC-seq fragment size distributions for mock and HSV-1 nuclei demonstrating expected nucleosomal banding patterns, including nucleosome-free (∼60 bp) and mono-nucleosome (∼200 bp) fragments.

**E)** Transcription start site (TSS) enrichment profiles for mock and HSV-1 samples. Aggregated insertion signal is plotted relative to annotated TSSs (±1 kb).

**F)** Cell state composition of single nuclei following clustering and annotation. Bar plots show the number of nuclei per annotated cell state in mock and HSV-1 conditions.

**Supplementary Figure 4. Extended SCENIC+ regulon activity and regulon size distributions across annotated brain cell populations**

**A)** Extended regulon activity across annotated cell populations. Heatmap of extended TF regulon activity across all annotated cell populations. Rows are cell populations and columns are extended TF regulons inferred by SCENIC+. Values indicate z-scored AUCell enrichment scores calculated per population. Hierarchical clustering was performed on both axes.

**B)** Distribution of regulon sizes for direct and extended SCENIC+ models. The number of predicted target genes per transcription factor regulon inferred using direct (motif-supported) and extended SCENIC+ models across all annotated cell populations. The distribution of regulon sizes is comparable between inference modes (median targets: Direct = 71; Extended = 52.5).

## Supplementary Data

Supplementary Data 1. Full direct and extended SCENIC+ regulon target lists.

Supplementary Data 2. Differential regulon activity statistics (ΔAUCell, p-values, adjusted p-values) for IFN-responsive vs homeostatic microglia and HSV-1 vs mock infiltrating macrophages.

Supplementary Data 3. Extended regulon connectivity table listing transcription factors and their inferred target genes.

Supplementary Data 4. Transcription factor–level summary metrics (ΔAUCell and total inferred target connectivity) used for Figure 4C and 4D analyses.

## Notes

### Competing Interest Statement

The authors have declared no competing interest.

### Summary of Updates

author affiliations and minor edits to text have been added.

