## Supplementary figures and images for "Intranasal HSV-1 Infection Drives Region-Specific Interferon-Dominant Microglial Remodeling"

### Supplemental Figure 1A

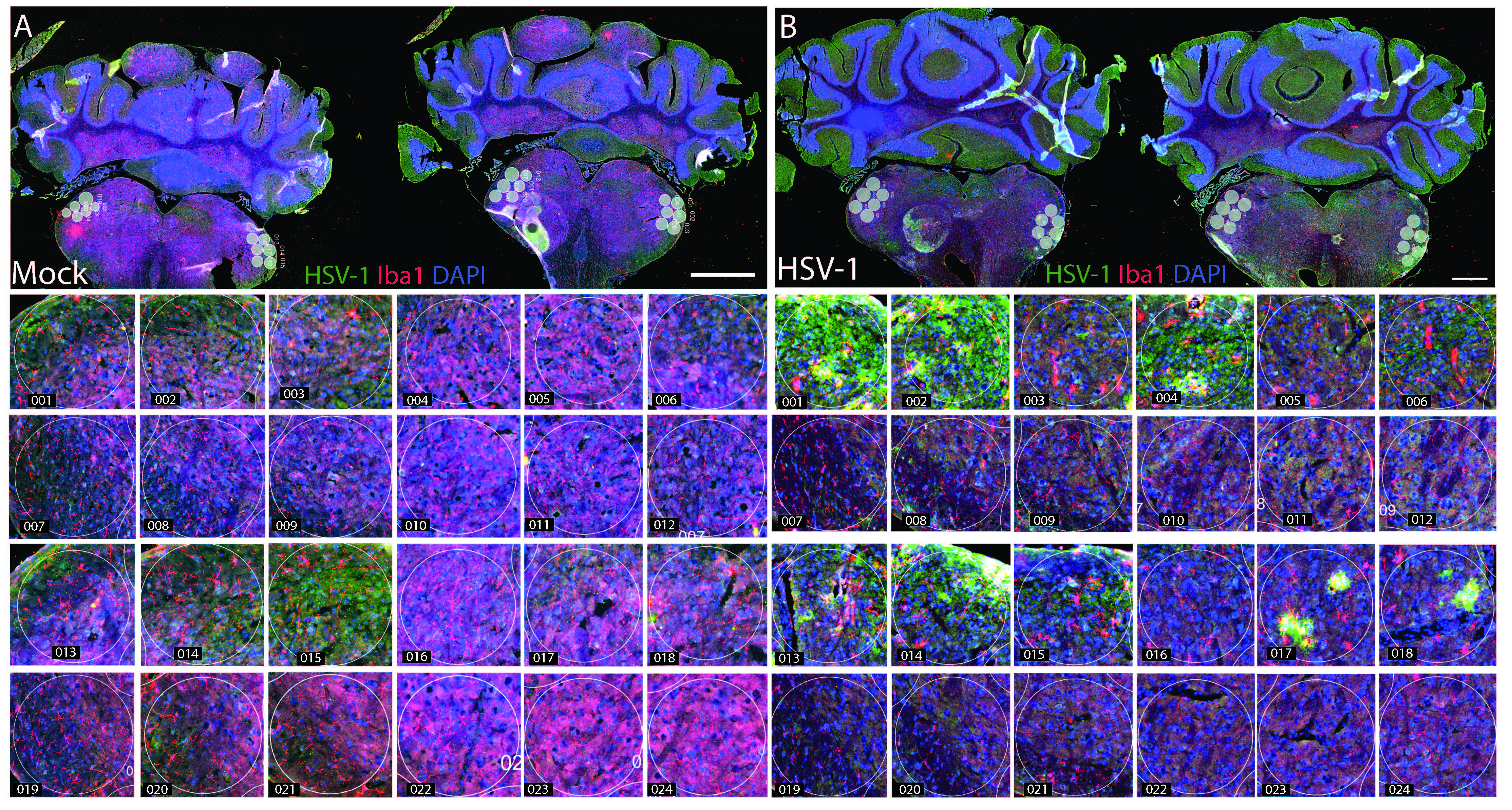

### Supplemental Figure 1B

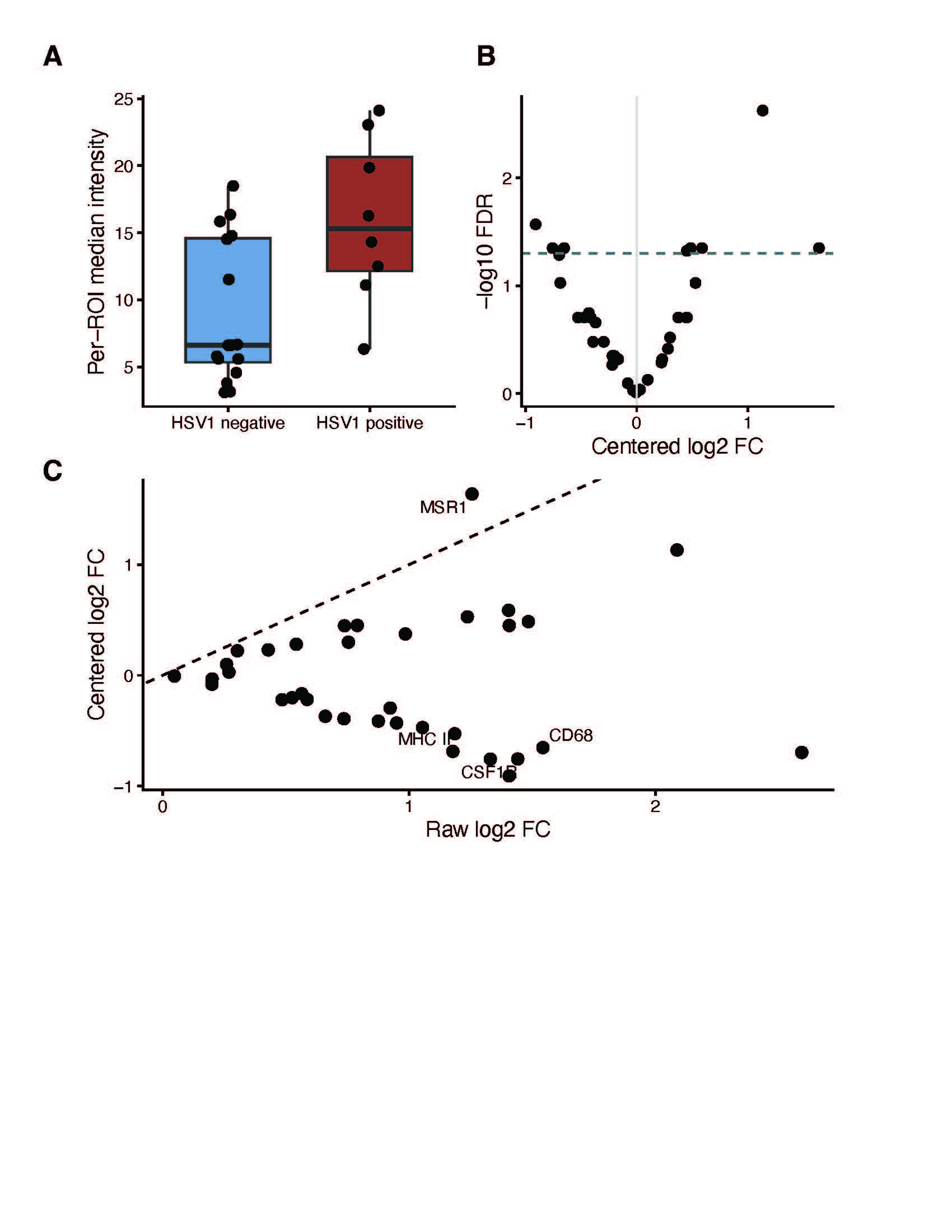

### Supplemental Figure 2

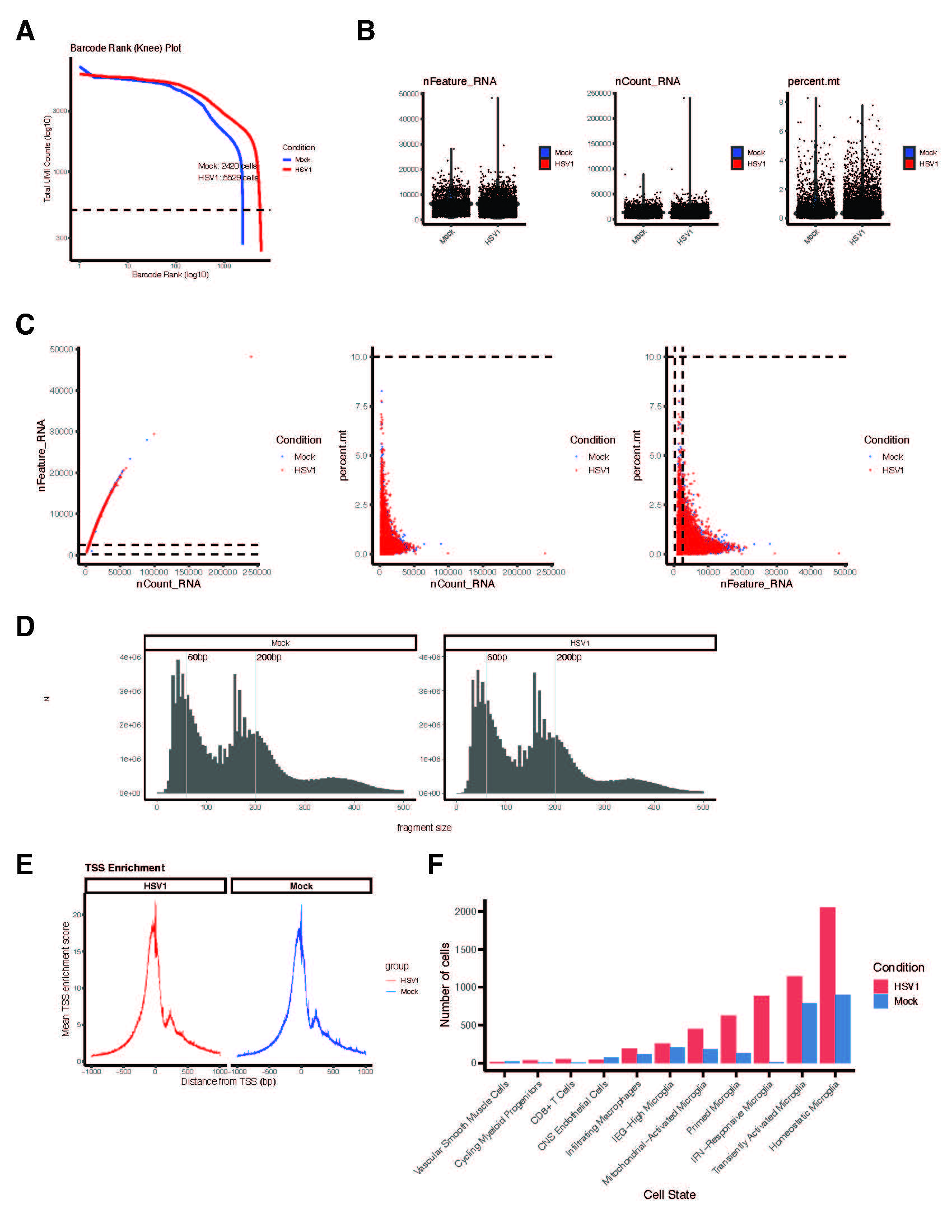

### Supplemental Figure 4

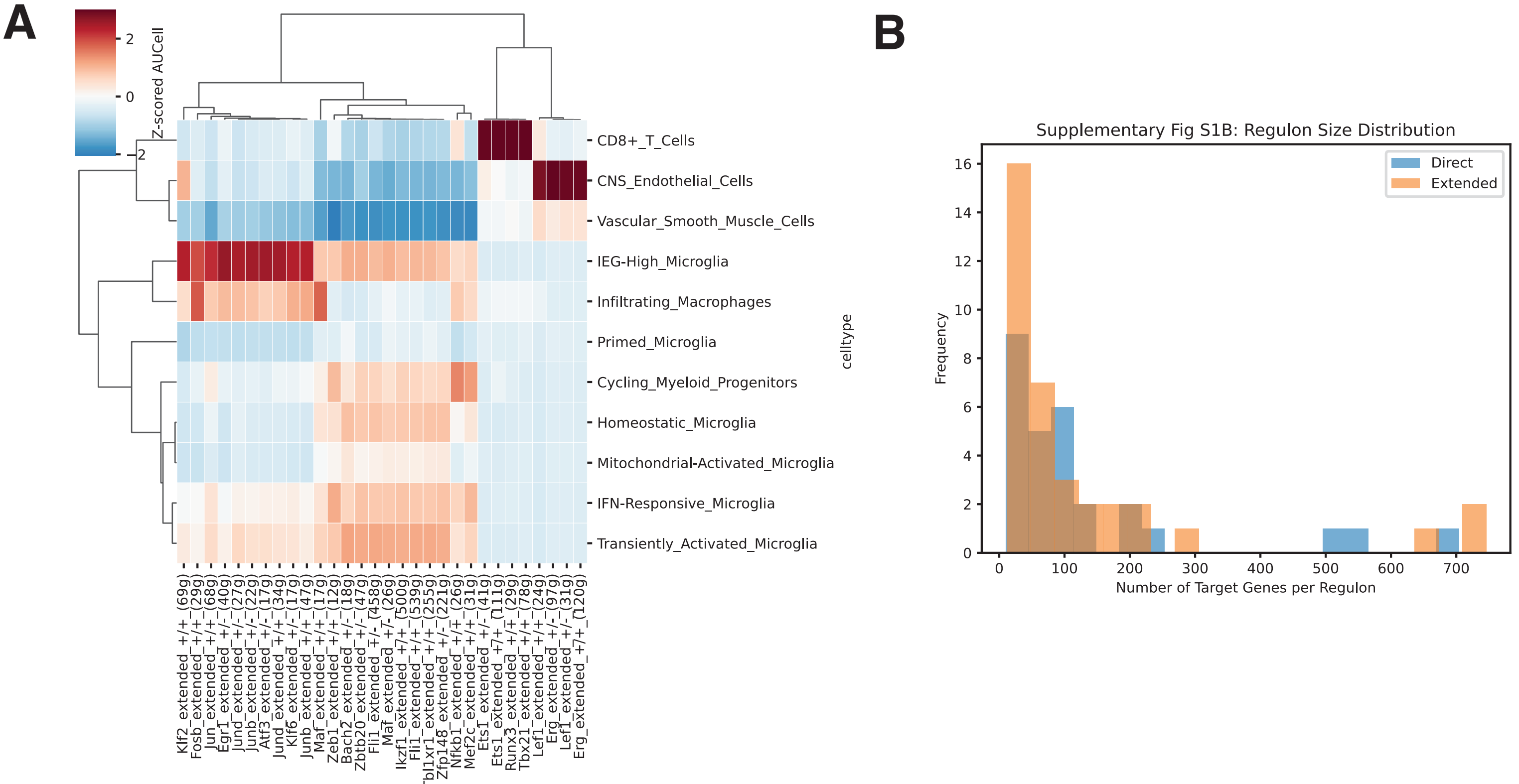
